# Overview of the SARS-CoV-2 genotypes circulating in Latin America during 2021

**DOI:** 10.1101/2022.08.19.504579

**Authors:** Jose Arturo Molina-Mora, Jhonnatan Reales-González, Erwin Camacho, Francisco Duarte-Martínez, Pablo Tsukayama, Claudio Soto-Garita, Hebleen Brenes, Estela Cordero-Laurent, Andrea Ribeiro dos Santos, Cláudio Guedes Salgado, Caio Santos Silva, Jorge Santana de Souza, Gisele Nunes, Tatiane Negri, Amanda Vidal, Renato Oliveira, Guilherme Oliveira, José Esteban Muñoz-Medina, Angel Gustavo Salas Lais, Guadalupe Mireles-Rivera, Ezequiel Sosa, Adrián Turjanski, María Cecilia Monzani, Mauricio G. Carobene, Federico Remes Lenicov, Gustavo Schottlender, Darío A. Fernández Do Porto, Jan Frederik Kreuze, Luisa Sacristán, Marcela Guevara-Suarez, Marco Cristancho, Rebeca Campos-Sánchez, Alfredo Herrera-Estrella

## Abstract

Latin America is one of the regions in which the COVID-19 pandemic has had a stronger impact, with more than 72 million reported infections and 1.6 million deaths until June 2022. Since this region is ecologically diverse and is affected by enormous social inequalities, efforts to identify genomic patterns of the circulating SARS-CoV-2 genotypes are necessary for the suitable management of the pandemic.

To contribute to the genomic surveillance of the SARS-CoV-2 in Latin America, we extended the number of SARS-CoV-2 genomes available from the region by sequencing and analyzing the viral genome from COVID-19 patients from seven countries (Argentina, Brazil, Costa Rica, Colombia, Mexico, Bolivia and Peru). Subsequently, we analyzed the genomes circulating mainly during 2021 including records from GISAID database from Latin America.

A total of 1534 genome sequences were generated from seven countries, demonstrating the laboratory and bioinformatics capabilities for genomic surveillance of pathogens that have been developed locally. For Latin America, patterns regarding several variants associated with multiple re-introductions, a relatively low percentage of sequenced samples, as well as an increment in the mutation frequency since the beginning of the pandemic, are in line with worldwide data. Besides, some variants of concern (VOC) and variants of interest (VOI) such as Gamma, Mu and Lambda, and at least 83 other lineages have predominated locally with a country-specific enrichments.

This work has contributed to the understanding of the dynamics of the pandemic in Latin America as part of the local and international efforts to achieve timely genomic surveillance of SARS-CoV-2.

## Introduction

In December 2019, several cases of a new respiratory illness were described in Wuhan, China. About a month later, it was confirmed that the illness COVID-19 (coronavirus disease 2019) was caused by a novel coronavirus which was subsequently named SARS-CoV-2 (Ashour, Elkhatib, Rahman, & Elshabrawy, 2020; World Health Organization, 2020). Until June 2022, the COVID-19 pandemic has impacted the world with >549 million confirmed cases of COVID-19, including >6.3 million deaths. Latin America is one of the most strongly impacted regions with more than 72 million reported infections and >1.6 million deaths during the same period.

SARS-CoV-2 genome sequences have been reported from many regions of the world and these data have been proven useful in tracking the global spread of the virus. Genomic epidemiology of SARS-CoV-2 has shed light on the origins of regional outbreaks, global dispersal, and epidemiological history of the virus (Gouvêa dos Santos, 2021; Young et al., 2020). Until April 2022, over 11.5 million genomes had been deposited in the GISAID database (https://www.gisaid.org/), out of which >376 000 were reported by Latin American countries.

Since its appearance, a large genetic diversity has been recognized for SARS-CoV-2 due to widespread transmission and geographical isolation (Zhou et al., 2021). The emergence of new genotypes (lineages, clades, variants, etc.) is the product of a natural process that occurs when viruses replicate at high rates as it happens during a pandemic (Gouvêa dos Santos, 2021). The World Health Organization (WHO) has classified five divergent genotypes as variants of concern (VOC: Alpha, Beta, Gamma, Delta, Omicron), as well as some lineages into variants of interest (VOI: Lambda, Mu, Epsilon, Zeta, Theta, Iota, Eta, Kappa, and others) and variants under monitoring (VUM: B.1.640 and XD) (World Health Organization, 2022). All the reported variants and other lineages have been identified in Latin America (Pan American Health Organization, 2021), including genotypes that were first reported regionally, such as Gamma in Brazil, Mu in Colombia, and Lambda in Peru (World Health Organization, 2022), as well as unique lineages in Costa Rica and Central America (Molina-Mora, 2022; Molina-Mora et al., 2021). Those descriptions of locally enriched genotypes exemplify the opportunities that SARS-CoV-2 has found in Latin America for spreading and evolving. This scenario is in part explained by the complex environmental and human reality in this region, with huge ecological diversity and social inequalities (Callejas, Echevarría, Carrero, Rodríguez-Morales, & Moreira, 2020; The Lancet, 2020). Thus, efforts on revealing the behavior of SARS-CoV-2 are necessary to identify regionally emerging patterns for the suitable management of the pandemic, which cannot be inferred from North America, Europe, or Asia (Callejas et al., 2020).

In this context, the CABANA initiative (Capacity building for Bioinformatics in Latin America, Global Challenges Research Fund GCRF: www.cabana.online) supported the development of a regional project titled “The SARS-CoV-2 genome, its evolution and epidemiology in Latin America” during 2021. The project had the direct participation of seven institutions from Argentina, Brazil, Bolivia, Colombia, Costa Rica, Mexico, and Peru. Efforts of this project included not only the sequencing and genome assembly of the SARS-CoV-2 virus from a total of 1534 COVID-19 cases in those countries, but also to bring a more complete overview of the SARS-CoV-2 genotypes circulating in Latin America during 2021 using public databases. Thus, this study aimed to contribute to the genomic surveillance of the SARS-CoV-2 to understand the dynamics of the pandemic in Latin America by providing genome sequences and analyzing circulating genotypes during the year 2021.

## Methods

### Samples and Ethical Considerations

Respiratory samples were obtained from public and private laboratories belonging to the national network of SARS-CoV-2 diagnostics in each country. Adequate transportation and storage conditions were guaranteed to preserve the samples. Every sample was anonymized to protect patients’ identity. Being a notifiable disease, the metadata was collected from the forms that accompanied the samples, either in the national reference laboratories or in the ministries of health. See Supplementary file for IDs to access metadata in the GISAID database.

### Sample sequencing and genome analysis

To contribute with SARS-CoV-2 genome sequences from Latin America, seven participant countries (Argentina, Bolivia, Brazil, Costa Rica, Colombia, Mexico, and Peru) were involved in sample processing from COVID-19 patients. Diagnosis using RT-qPCR, genome sequencing and assembly, as well as the genotyping, were implemented using the laboratory protocols and bioinformatic pipelines that are being locally used as part of the genomic surveillance efforts in each country as shown in Table 1 and reported in (Molina-Mora et al., 2021; Oliveira et al., 2022; Resende et al., 2020). Genome sequences were uploaded to the GISAID database (https://www.gisaid.org/). Details regarding the number of processed samples (assembled genomes), protocols, and GISAID accession numbers (ID) for each country are summarized in Table 1.

**Table 1.** Sequencing strategy and bioinformatic pipelines used for the genomic surveillance of the SARS-CoV-2 in five Latin American countries, CABANA initiative.

| **Country** | **Number of samples** | **Participant institutions** | **Sequencing protocol** | **Bioinformatic pipeline** |
| --- | --- | --- | --- | --- |
| Argentina | 220 | Universidad de Buenos Aires (UBA) & Consejo Nacional de Investigaciones Científicas y Técnicas (CONICET) | Illumina platform, following a custom protocol. | Genome assembly, variant calling, and genotyping: Custom protocol. |
| Bolivia | 94 | Laboratorio de diagnóstico e investigación BIOSCIENCE SRL & Laboratorio de Genómica Microbiana, Universidad Peruana Cayetano Heredia | Illumina platform, following a custom protocol. | Genome assembly, variant calling, and genotyping: Custom protocol. |
| Brazil | 167 | Vale Institute of Technology, Belém, PA, Brazil | Illumina platform, following a custom protocol. | Genome assembly, variant calling, and genotyping (Oliveira et al., 2022) |
| Costa Rica | 190 | Universidad de Costa Rica (UCR) & Instituto Costarricense de Investigación y Enseñanza en Nutrición y Salud (INCIENSA) | Illumina platform, following the protocol described in (Molina-Mora et al., 2021; Resende et al., 2020) | Genome assembly, variant calling, genotyping, and phylogeny: (Molina-Mora et al., 2021) |
| Colombia | 147 | Universidad de Los Andes. | Nanopore platform, following a custom protocol. | Genome assembly: custom protocol for long-reads sequencing.<br>Variant calling, genotyping, and phylogeny: (Molina-Mora et al., 2021) |
| Mexico | 472 | Unidad de Genómica Avanzada del Centro de Investigación y de Estudios Avanzados (UGA-LANGEBIO, CINVESTAV) | Illumina platform, following a custom protocol. | Genome assembly, variant calling, genotyping, and phylogeny: (Molina-Mora et al., 2021) |
| Peru | 244 | Laboratorio de Genómica Microbiana, Universidad Peruana Cayetano Heredia | Illumina platform, following a custom protocol. | Genome assembly, variant calling, and genotyping: Custom protocol. |

### Analysis of circulating SARS-CoV-2 genotypes in Latin America

To gain insights into the SARS-CoV-2 genotypes circulating in Latin America during 2021, a general analysis was done using the genome sequences available at the GISAID database (https://www.gisaid.org/). Selection of countries, statistics of sequenced samples, and plots of circulating genomes and mutation frequency were obtained using the tools of the GISAID platform. The number of COVID-19 cases per country was retrieved from the daily reports of the Pan American Health Organization (Pan American Health Organization, 2022). All the analyses were performed considering sequences collected until January 31^th^ 2022. PANGOLIN lineage database (O’Toole et al., 2021; Rambaut et al., 2020) was used to analyze the frequency of lineages among countries.

## Results and Discussion

Genomic surveillance has been a hallmark of the COVID-19 pandemic that, in contrast to other pandemics, achieves the tracking of the virus evolution and spread worldwide almost in real-time (Gouvêa dos Santos, 2021).

In this work, we extended the repertoire of SARS-CoV-2 genome sequences with a total of 1534 sequences from seven Latin American countries (Table 1). Whereas, this was a relatively modest contribution to the overall quantity of sequences produced in this period in Latin America for certain time-intervals and countries it provided important complementarity for the genomic surveillance of the virus. In Bolivia for example, our efforts represented 38% of all sequences produced over this time. To perform a more complete examination, we included all sequences from Latin America available at the GISAID database collected from February 2021 to January 2022.

According to the GISAID database records, the numbers of sequences is still small in comparison to the number of diagnosed cases in Latin America (Table 2). On average, only 0.39% of COVID-19 cases in Latin America have been sequenced, with Mexico and Chile having the highest rates with 0.98% and 0.92% respectively. In the case of Nicaragua, in which the pandemic has been downplayed (Pearson, Prado, & Colburn, 2020; Salazar Mather et al., 2020), the reports of diagnosed patients and other statistics are considered unrealistic, including the 2.92% of sequenced samples. Thus, we did not conduct comparisons of Nicaragua among other countries due to the extremely biased data.

**Table 2.** Comparison of COVID-19 cases and sequenced samples among Latin American countries until January 31th 2022.

| Country | Population | Total COVID-19 cases |  |  | Sequenced samples |  |
| --- | --- | --- | --- | --- | --- | --- |
|  |  | Absolute number | Percentage of the population (%) | Deaths | Absolute number | Percentage of COVID-19 cases (%) |
| Mexico | 131026542 | 4942590 | 3.77 | 306091 | 48329 | **0.98** |
| Chile | 19369864 | 2190561 | 11.31 | 39733 | 20086 | **0.92** |
| Belize | 408778 | 50487 | 12.35 | 625 | 376 | **0.74** |
| Ecuador | 18057002 | 732038 | 4.05 | 34533 | 3770 | **0.52** |
| Peru | 33681601 | 3239538 | 9.62 | 205834 | 14790 | **0.46** |
| Brazil | 214895351 | 25454105 | 11.84 | 627589 | 101532 | **0.40** |
| Costa Rica | 5166024 | 694865 | 13.45 | 7575 | 2719 | **0.39** |
| El Salvador | 6536807 | 135109 | 2.07 | 3899 | 308 | **0.23** |
| Colombia | 51721161 | 5887261 | 11.38 | 13400 | 12520 | **0.21** |
| Panama | 4419710 | 700274 | 15.84 | 7732 | 1223 | **0.17** |
| Guatemala | 18427485 | 690290 | 3.75 | 16385 | 1420 | **0.21** |
| Paraguay | 7267941 | 583662 | 8.03 | 17321 | 882 | **0.15** |
| Uruguay | 3492345 | 668425 | 19.14 | 6479 | 717 | **0.11** |
| Argentina | 45836859 | 8378656 | 18.28 | 121273 | 11553 | **0.14** |
| Honduras | 10147994 | 391874 | 3.86 | 10504 | 116 | **0.03** |
| Venezuela | 28311334 | 485974 | 1.72 | 5447 | 123 | **0.03** |
| Bolivia | 11919079 | 855705 | 7.18 | 20951 | 248 | **0.03** |
| Nicaragua | 6746365 | 17650 | 0.26 | 216 | 516 | **2.92** |
| Latin America | 617432242 | 56099064 | 9.09 | 1445587 | 221228 | **0.39** |
| Worldwide | 7900000000 | 380099991 | 4.81 | 5695345 | 7748697 | **2.04** |

On the other extreme, Bolivia, Honduras and Venezuela have barely sequenced even 0.03% of samples derived from all patients diagnosed with the disease. There is no single Latin American country that has sequenced more samples, relative to the number of cases reported, than the world average that corresponds to 2.04%, which is low too. The current scenario is congruent with a previous report with less than 0.5% of sequenced samples for Latin American countries (Brito et al., 2021). These findings represent not only part of the regional disparities in the SARS-CoV-2 genomic surveillance efforts in Latin America, but also that this geographic region needs to increase the effort to achieve the sequencing of at least 5% of positive samples to detect emerging viral lineages when their prevalence is less than 1% of all strains in a population, as suggested previously (Vavrek et al., 2021). This situation is like that of other latitudes around the world in which only a very small portion of the countries has reached the recommended percentage, suggesting that sequencing at least 0.5% of the cases, with a time in days between sample collection and genome submission <21 days, could be a benchmark for SARS-CoV-2 genomic surveillance efforts for low- and middle-income countries (Brito et al., 2021) taking into account the high cost of sequencing reagents and equipment in these countries. Thus, the identification of patterns regarding the circulating genotypes in Latin America should be interpreted with cautions due the differences of SARS-CoV-2 surveillance systems, including sequencing capacity and sampling strategies between countries in the region.

Regarding the circulating genotypes, the reports on the diversity of lineages are similar to other studies in Latin America (Laiton-Donato et al., 2021; Molina-Mora et al., 2021; Romero et al., 2021; Vargas-Herrera et al., 2022) and other distant geographic regions (Alizon et al., 2021; Wang et al., 2021). For divergent SARS-CoV-2 genomes, all VOCs have been reported in all Latin American countries, resulting in a large diversity of genotypes circulating in each country (Figure 1). This is in line with the expected pattern of multiple and independent re-introductions due to population mobility within Latin America, as well as to and from other countries and continents (da Silva Filipe et al., 2020; Ramírez et al., 2021; Spanakis et al., 2021). Besides, some genotypes have been reported with an epicenter in Latin America. As presented in Figure 2, those country-specific variants were predominant in the first semester of 2021, such as the Gamma variant in Brazil until August 2021, the Mu variant in Colombia from April to September 2021, and Lambda in Peru during the period from March to June 2021 (GISAID, 2021). Other remarkable genome versions were the case of the Gamma variant predominating between June and August 2021 in Argentina, as well as the more mixed pattern with distinct variants in Mexico, similar to the average for the entire Latin American region. In comparison to the rest of the world, Latin America reported similar transitions between the Alpha, Delta, and Omicron variants. Nonetheless, the increased reports of Mu and Lambda in this region were minimal for the worldwide representation (Figure 2).

**Figure 1.**
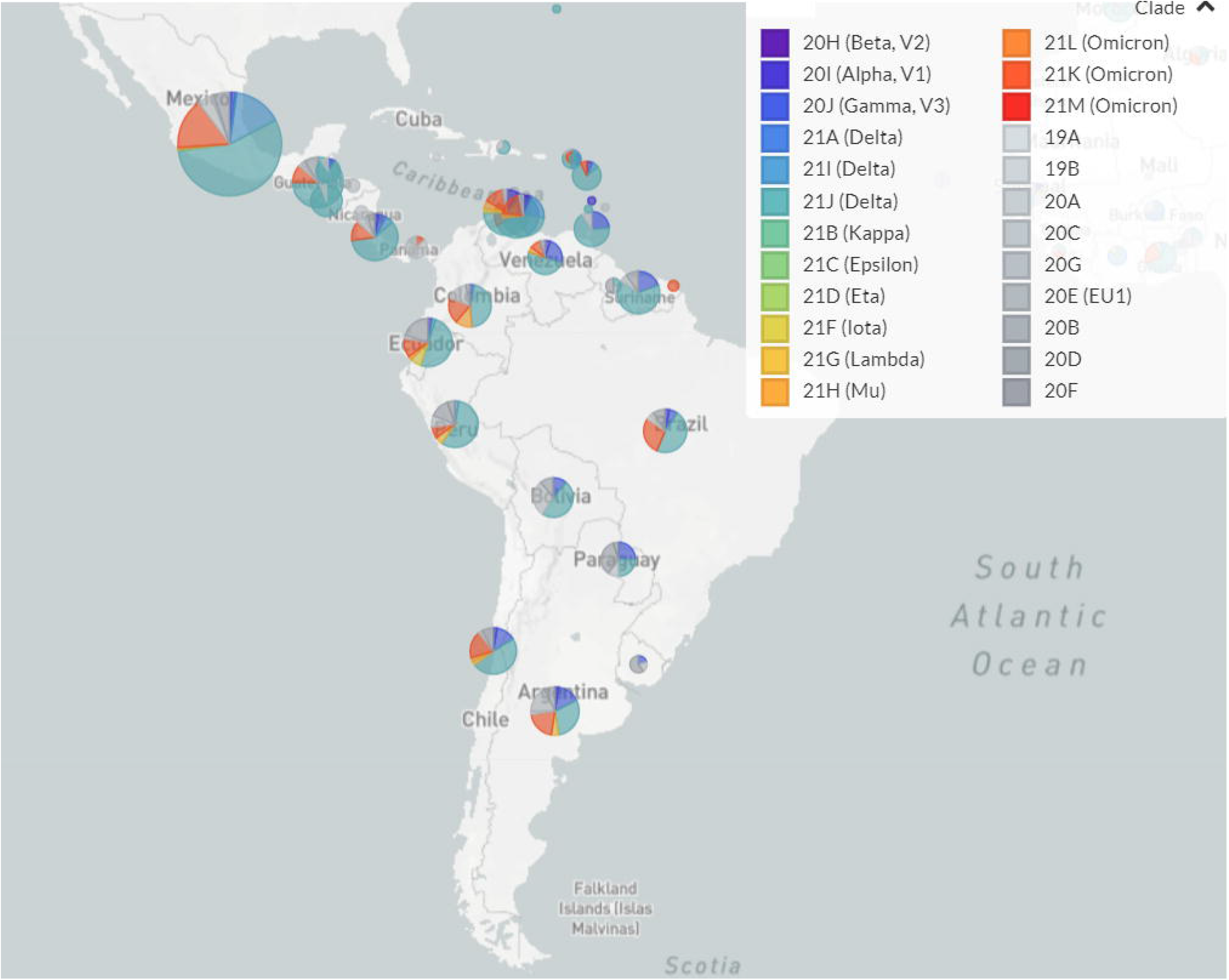
Landscape of the SARS-CoV-2 genotypes circulating in Latin America until January 2022. Pie charts indicate the relative abundance of distinct SARS-CoV-2 genotypes in each country.

**Figure 2.**
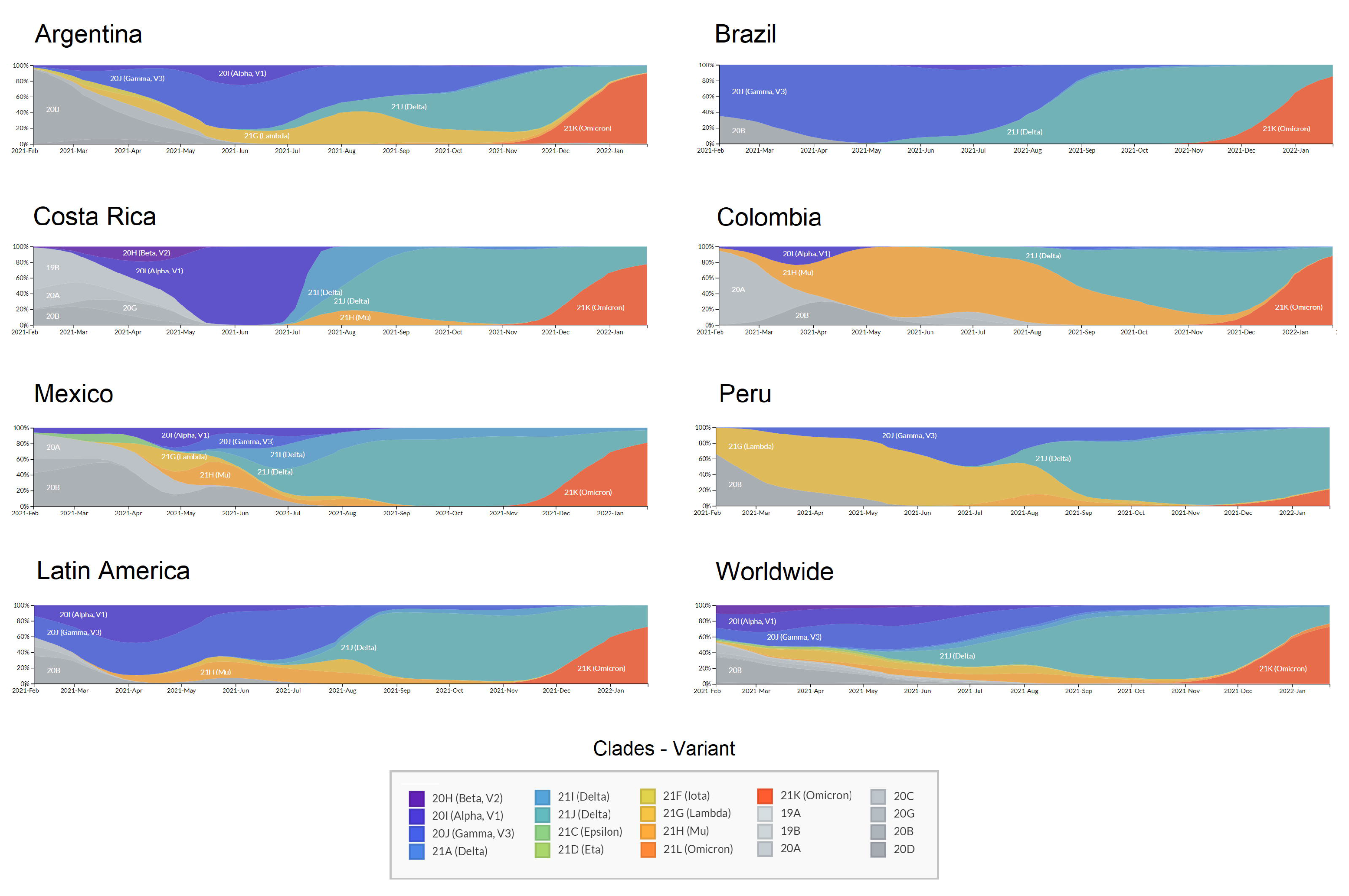
Transition of SARS-CoV-2 variants circulating in six Latin American countries and worldwide during 2021. X: time from February 2021 to January 2022; Y: Relative abundance of SARS-CoV-2 genotypes.

In Latin America recurrent dissemination of SARS-CoV-2 through shared borders between countries has been evidenced (Paniz-Mondolfi et al., 2020), allowing rapid entrance and dissemination of different lineages to the different countries (Mir et al., 2021). Territories with no restriction to international interchange are more likely to introduce multiple SARS-CoV-2 variants, including variants of concern and/or interest and even lineages with mutations of concern and emerging variants with different mutation patterns (Williams et al., 2021). These introductions of VOCs to Latin America were more evident during the second half of the year 2021, where the Delta variant displaced other variants in several countries and became predominant as shown in Figure 2, while during the first semester of the year lineage predominance varied among these countries.

Although several epidemiological aspects can be associated with these patterns, the extensive opening of the borders during the middle of 2021 possibly favored the spread of new variants of concern in the region. Besides, the presence of multiple mutations that have been associated with increased infectivity and/or escape from immune response in variants such as Delta (Shiehzadegan, Alaghemand, Fox, & Venketaraman, 2021) helped this variant to displace other variants, as it occurred worldwide.

For other genotypes, at least 83 out of >1500 PANGOLIN lineages have been reported with a high predominance in a country Latin American country (Table 3). The full list of lineages is presented in the supplementary file. As an example, lineage C.39 was predominant in Chile with 45.0% of all the sequences reported, followed by France (15.0%), Peru (12.0%), Guinea (10.0%), and Germany (8.0%). From these lineages, at least 80% of the sequences from 51 lineages have been reported to come from a Latin American country (Table 3 and Table 4). In the distribution by country, Brazil, Peru, Chile, Costa Rica, and Mexico have more reports of lineages with a frequency >80% locally.

**Table 3.** Number of lineages in which a Latin American country is predominant by frequency.

| **Country** | **Lineages in which the country is predominant\*** | **Lineages in which the country has frequency >80%\*\*** |
| --- | --- | --- |
| Argentina | 4 | 3 |
| Brazil | 30 | 21 |
| Chile | 11 | 6 |
| Colombia | 2 | 0 |
| Costa Rica | 6 | 3 |
| Ecuador | 1 | 0 |
| Mexico | 5 | 2 |
| Panama | 1 | 1 |
| Peru | 21 | 13 |
| Uruguay | 2 | 2 |
| Total | 83 | 51 |

\* Refers to those lineages in which the main source of sequences is from a Latin American country. See text for details.
\*\* Similar to the previous case, but lineages are only counted if the predominant country has a percentage >80%. See text for details.

**Table 4.** Lineages reported with a frequency >80% among Latin American countries.

| Lineage | Most common countries | First detected - Country | Total Sequences worldwide |
| --- | --- | --- | --- |
| A.2.4 | Panama 96.0%, Costa_Rica 2.0%, Northern_Mariana_Islands 1.0%, United States of America 1.0%, United Kingdom 0.0% | Panama | 291 |
| C.4 | Peru 95.0%, United States of America 2.0%, Turkey 2.0%, Republic of Serbia 1.0%, Switzerland 1.0% | Peru | 130 |
| C.11 | Chile 97.0%, Peru 2.0%, Argentina 1.0% | Chile | 95 |
| C.13 | Peru 84.0%, United States of America 14.0%, Japan 2.0% | Peru | 45 |
| C.14 | Peru 92.0%, United States of America 3.0%, Japan 2.0%, Democratic Republic of the Congo 1.0%, Brazil 1.0% | Peru | 253 |
| C.25 | Peru 100.0% | Peru | 8 |
| C.29 | Chile 89.0%, Russia 6.0%, Denmark 6.0% | Chile | 18 |
| C.32 | Peru 95.0%, Turkey 2.0%, Denmark 2.0%, United Kingdom 2.0% | Peru | 57 |
| C.33 | Peru 90.0%, Japan 5.0%, United States of America 5.0% | Peru | 20 |
| C.40 | Peru 97.0%, Chile 3.0% | Peru | 37 |
| P.1.3 | Brazil 100.0% | Brazil | 29 |
| P.1.4 | Brazil 98.0%, Costa_Rica 1.0%, Peru 1.0%, United States of America 0.0%, Colombia 0.0% | Brazil | 1039 |
| P.1.5 | Brazil 100.0% | Brazil | 14 |
| P.1.6 | Brazil 99.0%, Sweden 0.0%, French_Guiana 0.0% | Brazil | 562 |
| P.1.7 | Brazil 91.0%, United States of America 5.0%, Spain 2.0%, Peru 0.0%, Mexico 0.0% | Japan | 3666 |
| P.1.7.1 | Peru 94.0%, United States of America 2.0%, Brazil 1.0%, Chile 1.0%, Puerto_Rico 0.0% | Bolivia | 773 |
| P.1.8 | Brazil 94.0%, Sweden 1.0%, France 1.0%, United Kingdom 1.0%, Spain 1.0% | Brazil | 257 |
| P.1.9 | Brazil 97.0%, United States of America 1.0%, Mexico 1.0%, Belgium 1.0% | Brazil | 236 |
| P.1.10.2 | Mexico 100.0% | Mexico | 23 |
| P.1.11 | Brazil 96.0%, Ecuador 2.0%, Spain 1.0%, United States of America 1.0% | Brazil | 102 |
| P.1.12.1 | Peru 91.0%, United States of America 4.0%, Italy 2.0%, Chile 1.0%, Switzerland 1.0% | Italy | 161 |
| P.4 | Brazil 100.0%, United States of America 0.0% | Brazil | 234 |
| P.5 | Brazil 95.0%, Philippines 3.0%, United States of America 2.0% | Brazil | 44 |
| P.6 | Uruguay 95.0%, Philippines 2.0%, United States of America 1.0%, Norway 0.0%, Spain 0.0% | Uruguay | 298 |
| B.1.1.33 | Brazil 83.0%, United States of America 5.0%, Chile 3.0%, Argentina 2.0%, Paraguay 1.0% | United States | 2130 |
| N.3 | Argentina 97.0%, Bolivia 2.0%, Hong_Kong 1.0%, Chile 1.0% | Argentina | 124 |
| N.4 | Chile 92.0%, Brazil 4.0%, United States of America 1.0%, Peru 1.0%, New_Zealand 0.0% | Canada | 232 |
| N.5 | Argentina 87.0%, United States of America 7.0%, Spain 1.0%, Italy 1.0%, United Kingdom 1.0% | India | 365 |
| N.6 | Chile 97.0%, Brazil 1.0%, Japan 1.0%, Paraguay 1.0% | Chile | 138 |
| N.7 | Uruguay 100.0% | Uruguay | 33 |
| N.9 | Brazil 96.0%, Ireland 1.0%, Argentina 1.0%, Japan 1.0%, Chile 1.0% | Brazil | 138 |
| N.10 | Brazil 97.0%, Germany 3.0% | Brazil | 22 |
| B.1.1.110 | Peru 92.0%, Finland 8.0% | Finland | 11 |
| B.1.1.324 | Chile 100.0% | Chile | 17 |
| B.1.1.332 | Brazil 100.0% | Brazil | 29 |
| B.1.1.389 | Costa_Rica 86.0%, United States of America 6.0%, Spain 5.0%, Finland 2.0%, Australia 1.0% | Costa Rica | 139 |
| B.1.1.442 | Argentina 80.0%, Turkey 7.0%, United States of America 5.0%, Germany 5.0%, Austria 2.0% | Argentina | 47 |
| B.1.1.516 | Costa_Rica 100.0% | Nicaragua | 21 |
| B.1.110.1 | Chile 82.0%, United States of America 18.0% | Chile | 12 |
| B.1.205 | Peru 98.0%, Israel 2.0% | Peru | 40 |
| B.1.243.2 | Mexico 81.0%, United States of America 19.0% | Mexico | 60 |
| B.1.291 | Costa_Rica 94.0%, United States of America 4.0%, Australia 2.0% | Nicaragua | 67 |
| AY.25.1.1 | Peru 82.0%, United States of America 14.0%, Chile 2.0%, Colombia 1.0%, Switzerland 1.0% | USA | 172 |
| AY.26.1 | Peru 81.0%, United States of America 12.0%, Mexico 2.0%, Israel 1.0%, Chile 1.0% | Peru | 154 |
| AY.43.1 | Brazil 96.0%, United Kingdom 1.0%, United States of America 1.0%, France 1.0%, Chile 1.0% | Poland | 1036 |
| AY.43.2 | Brazil 99.0%, France 0.0%, Japan 0.0%, Belgium 0.0%, Switzerland 0.0% | India | 1295 |
| AY.43.7 | Brazil 98.0%, Netherlands 1.0%, France 1.0%, Israel 1.0% | France | 184 |
| AY.46.3 | Brazil 95.0%, United States of America 1.0%, Turkey 1.0%, India 0.0%, Czech_Republic 0.0% | India | 1565 |
| AY.99.1 | Brazil 85.0%, United Kingdom 10.0%, United States of America 4.0%, India 1.0%, Spain 0.0% | India | 1372 |
| AY.99.2 | Brazil 97.0%, United States of America 1.0%, Chile 0.0%, France 0.0%, Portugal 0.0% | Brazil | 22815 |
| AY.101 | Brazil 81.0%, Chile 10.0%, Colombia 4.0%, Peru 2.0%, United States of America 1.0% | Colombia | 4300 |

For instance, Peru had 97.0% of the sequences reported for lineage C.40 and 95% of lineage C.4. Also, Peru was the main country in which the AY.25.1.1 and AY.26.1 genotypes (Delta sub-lineages) were documented. Brazil reported 97% of the 22 815 cases of lineage AY.99.2 (firstly reported in Colombia), that was demonstrated to successfully disseminate among different locations in the country (de Souza et al., 2022; Gularte et al., 2022). Lineages derived from the Gamma variant were also reported frequently in Brazil (e.g. P.1.4, P.1.7, P.4 and others). During 2020 the lineage B.1.1.389, which harbors the specific mutation spike:T1117, was reported as predominant in Costa Rica (86% of cases of this lineage were reported in this country) (Molina-Mora et al., 2021). Despite its dominance, few changes were predicted on the virus behavior (transmission, immune response, other) and it was quickly replaced by the lineage Central America and subsequently by VOCs such as Alpha and Gamma (Molina-Mora, 2022). In the case of Mexico, two lineages (P.1.10.2 and B.1.243.2) were mainly found in this country (frequency >80%) but in a limited number. Lineage B.1.1.519 was a relevant genotype reported in Mexico, despite it was mainly reported in the United States. This version predominated in Mexico during the first quarter of 2021 while the Alpha variant (B.1.1.7) was also spreading. Interestingly, unlike other cases, the Alpha variant did not displace B.1.1.519 in this country (Zárate et al., 2022). B.1.1.519 was assigned as a VUM by WHO in 02-Jun-2021 and was degraded to a FMV (formerly monitoring variant) on 9-Nov-2021 (https://www.who.int/activities/tracking-SARS-CoV-2-variants).

Jointly, these results indicate that specific mutations and the subsequent consolidation into lineages were detected in Latin America and evidenced by genomic surveillance in the region. Interestingly, 17 of these lineages were first reported in a different country from where it was subsequently found to predominant (>80%). This includes neighboring countries, such as the case of lineage P.1.7.1 which was enriched in Peru but was first reported in Bolivia. This pattern was more frequent for Brazil, with eight lineages that were first reported in other countries including from Europe and Asia, but that became dominant in this country.

Tracking of specific mutations into Latin American lineages that could be used as local markers, may help to identify transmission networks locally and globally, highlighting the need for each country and territory to strengthen the sequencing and bioinformatic capacities. These capacities can also be of use to locally study other scenarios such as clinical profiles for COVID-19 patients (Molina-Mora, González, et al., 2022), long-term COVID-19 (Reales Gonzalez et al., 2022), identification of co-infections (Molina-Mora, Cordero-Laurent, Calderón-Osorno, Chacón-Ramírez, & Duarte-Martínez, 2022) or identify recombinant genomes (a recognized mechanism of viral diversity in coronaviruses).

Despite the reports of differences in the enriched genotypes in the first half of 2021, the emergence of new variants of the viral genome in Latin America was consistent with the rest of the world inferred from the mutation frequency (Figure 3-A). During 2020, the mutation frequency for the S1 region of the spike gene was estimated at around 2-3 mutations per month. At the beginning of 2021, this frequency increased to 8.32 and subsequently to around 12 with the predominance of Delta. However, with the arrival of the Omicron variant, the frequency at the very end of 2021 and the first month of 2022 reached values of 28 mutations, in both, Latin America and the world. Thus, this accumulative divergence has impacted the mutation rate over the pandemic, which until January 2022 was estimated to be around 8.74 × 10^−4^ substitutions per site per year (Figure 3-B). This mutation frequency and rate values are consistent with other local and global studies during the pandemic (Bouckaert et al., 2019; Corey et al., 2021; Desai et al., 2021), including the rate of 0.8 - 2.38 × 10^−3^ substitutions per site per year described in (Banerjee, Mossman, & Grandvaux, 2021). Following the gradual reopening of borders and worldwide travels, the frequency of infections and the appearance of mutations and new genotypes are expected to increase (Liu et al., 2021). Thus, more genome sequencing studies, including robust metadata collection, and more financial support are needed to continue with the surveillance of the pandemic in Latin America.

**Figure 3.**
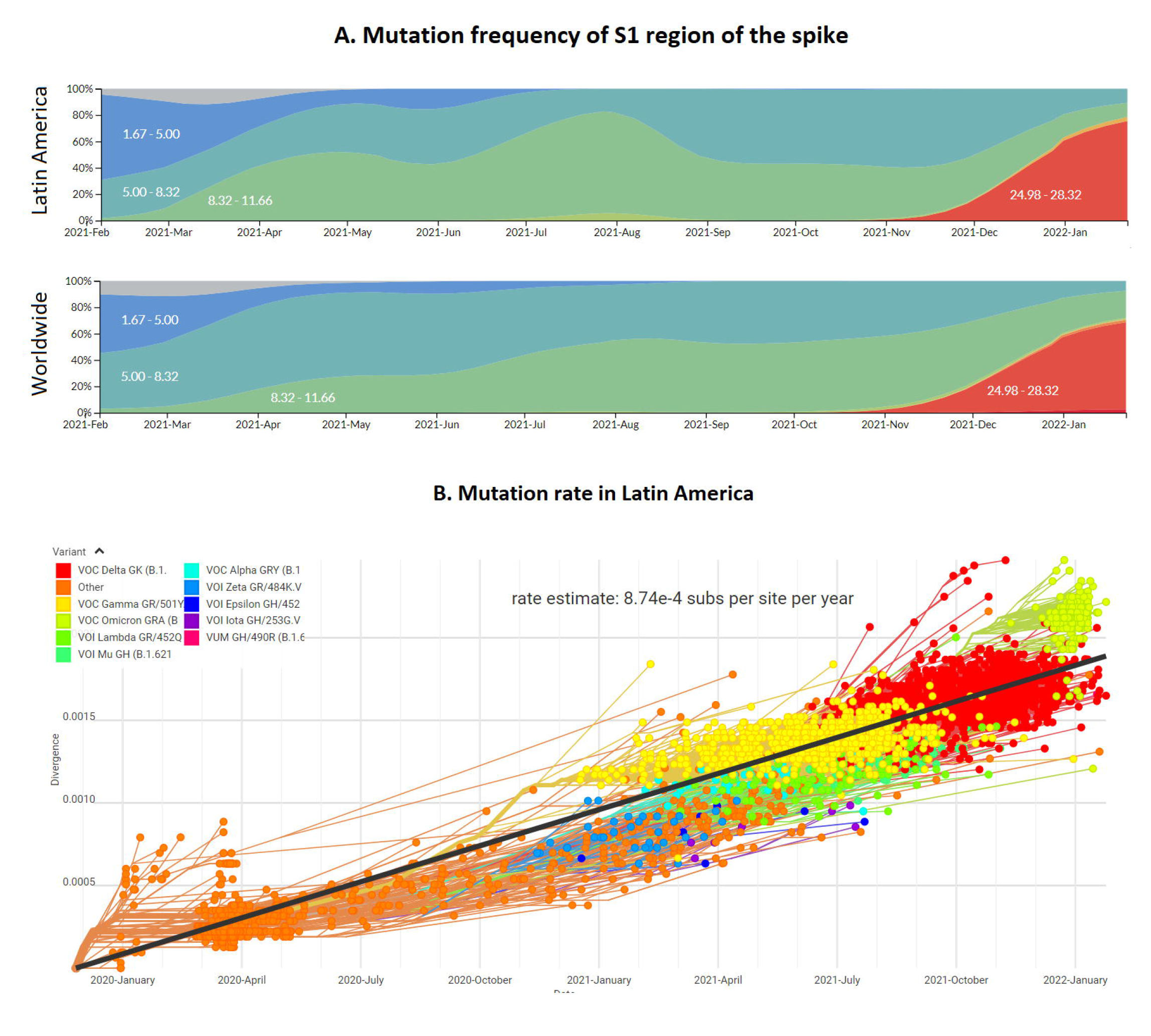
Mutation profile of the SARS-CoV-2 genome in Latin America and worldwide during 2021. (A) Mutation frequency in the region S1 encoding the spike protein of the SARS-CoV-2 viral genome. Colors represent the intervals for the absolute mutation number. (B) Mutation rate of the SARS-CoV-2 genomes during the pandemic, including distinct genotypes (colors) and the approach to estimate consensus rate (black line).

Finally, since most countries in this region are considered low- and middle-income countries, the impact of the COVID-19 pandemic on society has been devastating socially and economically (Bottan, Hoffmann, & Vera-Cossio, 2020). Genomic surveillance is pivotal as a powerful tool for decision-makers regarding the management of the pandemic in the Latin American context concerning social and economic measures, as well as practical decisions in terms of the diagnostic tools, treatments, and vaccines (Gouvêa dos Santos, 2021). On the other hand, local and prompt reports of emerging genotypes demonstrated the laboratory and bioinformatic capabilities in Latin American countries. These capabilities were developed locally in the last years for the surveillance of pathogens and other applications. Jointly, the local and international efforts to achieve the genomic surveillance of SARS-CoV-2 have contributed to the understanding of the dynamics of the pandemic in Latin America, which is an ongoing process.

## Conclusions

In conclusion, with this study we have contributed to the genomic surveillance of the SARS-CoV-2 in Latin America by providing 1534 genome sequences from seven countries and the subsequent global analysis of circulating genomes mainly during 2021. For Latin America, patterns regarding several variants associated with multiple re-introductions, a relatively low proportion of sequenced samples, as well as an increase in the mutation frequency, are in line with worldwide data. Additionally, some genotypes such as Gamma, Mu and Lambda variants and 83 lineages have emerged locally with a subsequent country-specific predominance. Regional efforts demonstrate the laboratory and bioinformatics capabilities for the genomic surveillance of pathogens that have been developed in Latin America, and which is expected to continue during the current COVID-19 pandemic.

## Acknowledgments

We thank to public and private clinical laboratories for the samples of confirmed cases of COVID-19 from the participant countries.

## Data availability

Raw data was uploaded to ENA under the project number PRJEB53987.

Assembled genomes and related metadata were uploaded to GISAID database; see accession numbers in supplementary file.

## Author contributions

J.A.M.M, P.T., G.O., J.F.K., M.C, R.C.S., and A.H.E participated in the conception and design of this study. F.D.M, C.S.G, H.B, E.C.L., J.E.M.M, A.V, A.R.S., C.G.S., C.S.S., J.S.S., A.G.S.L., L.S., M.G.S., and G.M.R. were involved in sample processing. J.A.M.M. standardized the bioinformatics pipelines. J.A.M.M., G.N., R.O., T.N., J.R.G, and E.C were involved in data analysis. J.A.M.M., F.D.M., O.G. J.F.K. M.C., R.C.S., and A.H.E. participated in the interpretation of results. J.A.M.M. drafted the manuscript. All authors reviewed and approved the final manuscript.

